# Single platelet variability governs population sensitivity and initiates intrinsic heterotypic behaviours

**DOI:** 10.1101/2020.01.22.915512

**Authors:** Maaike S. A. Jongen, Ben D. MacArthur, Nicola A. Englyst, Jonathan West

## Abstract

Droplet microfluidics combined with flow cytometry was used for high throughput single platelet function analysis. A large-scale sensitivity continuum was shown to be a general feature of human platelets from individual donors, with hypersensitive platelets coordinating significant sensitivity gains in bulk platelet populations and shown to direct aggregation in droplet-confined minimal platelet systems. Sensitivity gains scaled with agonist potency (convulxin>TRAP-14>ADP) and reduced the collagen and thrombin activation threshold required for platelet population polarization into pro-aggregatory and pro-coagulant states. The heterotypic platelet response results from an intrinsic behavioural program. The method and findings invite future discoveries into the nature of hypersensitive platelets and how community effects produce population level behaviours in health and disease.

## Introduction

Understanding cellular diversity and interactions provides the key to elucidating system behaviour. It becomes meaningful to investigate cellular diversity and identify even potentially rare phenotypes when amplification mechanisms exist in the system and when there is good reason to predict large-scale variety. Classically, cancer^1-3^, immunology^4,5^ and stem cells^6,7^ with associated cell expansion have been the focus of the large majority of single cell studies.

In this work we turn our attention to platelets, dispersed sentinels which patrol the vasculature to detect breaches and respond in a coordinated manner using rapid and potent paracrine signalling to collectively form a thrombus. Platelets are also inherently variable^8^, originating from the fragmentation of heterotypic^9^ megakaryocytes resulting in variously small sub-cellular compartments (60% volume CV)^10^ with dissimilar contents and biochemistry^11-13^ and, without a nucleus, in a state of decay^14,15^ before clearance. Therefore, platelet activation represents an ideal system for investigating cellular diversity and consequences for homeostatic system control. Indeed, the nature and functional consequences of platelet diversity has been a matter of enquiry for almost half a century^8,10,11^. More recently, the discovery that dual stimulation with collagen and thrombin^16,17^ polarises platelets into distinct pro-coagulant and pro-aggregatory phenotypes^8,18-24^ has renewed interest on the topic of platelet diversity. In particular, the pro-coagulant platelets have been further characterised^25-30^, revealing diverse functions that either represent multiple pro-coagulant subpopulations or a unified, yet versatile pro-coagulant subpopulation^21^. The bifurcation of the platelet population into the two phenotypes further creates debate regarding intrinsic versus extrinsic functional programming^8^. Allied to this, subjects with reduced GPVI levels showed reduced thrombus formation^31^, implicating platelet heterogeneity with increased activity by platelets with elevated GPVI levels^32^. Overall, a complex picture is emerging, with precision methods required to accurately delineate subpopulations and their functional roles to inform our understanding of platelet interactions governing thrombus formation.

The paracrine signalling inherent to platelet activation represents a technical challenge for measuring single platelet behaviour without interference by the secretion products of activated platelets in the vicinity. This implies the requirement for confinement, discretising the analysis into single platelet measurements. The other requirement is throughput to effectively resolve the functional structure of the platelet population. Droplet microfluidics allows the reliable production of monodisperse droplets in the nanolitre to femtolitre range and has emerged as a powerful tool for packaging single cells in high throughput^33-37^. Here, the surfactants assembled at the aqueous:oil interface prohibit exchange between other aqueous compartments to eliminate platelet–platelet cross-talk. Droplet-based analytical methods have been effectively applied to cell phenotyping^38-42^ and are also popularly used for single cell sequencing^43-46^. In this contribution we describe the first application of droplet microfluidics for mapping the functional behaviour of suspension platelet populations with single platelet resolution. Comparing the responses with bulk platelet populations demonstrates the existence of hypersensitive platelets which can coordinate system-level sensitivity gains, a feature shown to drive heterotypic system polarisation during dual agonist stimulation.

## Results and Discussion

Microfluidics is suited for the handling of blood cells that naturally exist in a suspension state. This is especially relevant for platelets, which are sufficiently small to be near-neutrally buoyant allowing sustained delivery to the microfluidic device without the need for stirring and associated shear effects^47^. Indeed, platelets are characteristically shear-sensitive^48,49^ and the droplet generation junction introduces shear conditions, albeit short-lived (∼50 μs). Critically, platelet activation was absent in the vehicle control samples demonstrating that the shear conditions for droplet generation, and droplet transport^50^, as well as the surfactant and fluorinated PDMS channel walls do not activate platelets.

The experimental concept is illustrated in Figure 1A along with consideration of the Poisson distribution in Figure 1B which informs the choice of droplet volume and/or platelet concentration required for the efficient encapsulation of single platelets. For a platelet concentration of 25 × 10^6^/mL and with further on-chip dilution (x5) with agonist and antibody volumes, this indicates that an 8 pL droplet volume (ø25 μm) produces effective single platelet encapsulation: 3.38% of droplets contain a single platelet, 0.08% contain multiples with a single to multiple ratio of 42. The droplet microfluidic circuit used in this study is shown in Figure 1C and was used to generate 25-μm-diameter droplets (Figure 1D,E) at 10.4 kHz for single platelet packaging (352 Hz). This allows >100,000 platelets to be encapsulated in the 5 minute collection timeframe. To demonstrate the Poisson distribution effect, high platelet input concentrations were used to observe the relationship between singlet and multiple occupancy events (Figure 1F). Coupled with the kHz measurement capabilities of flow cytometry the analytical pipeline enables the functional variety of large-scale platelet populations to be readily mapped. The complete sampling to microfluidics and flow cytometry methodology is illustrated in **Supplementary Figure 1**.

**Figure 1.**
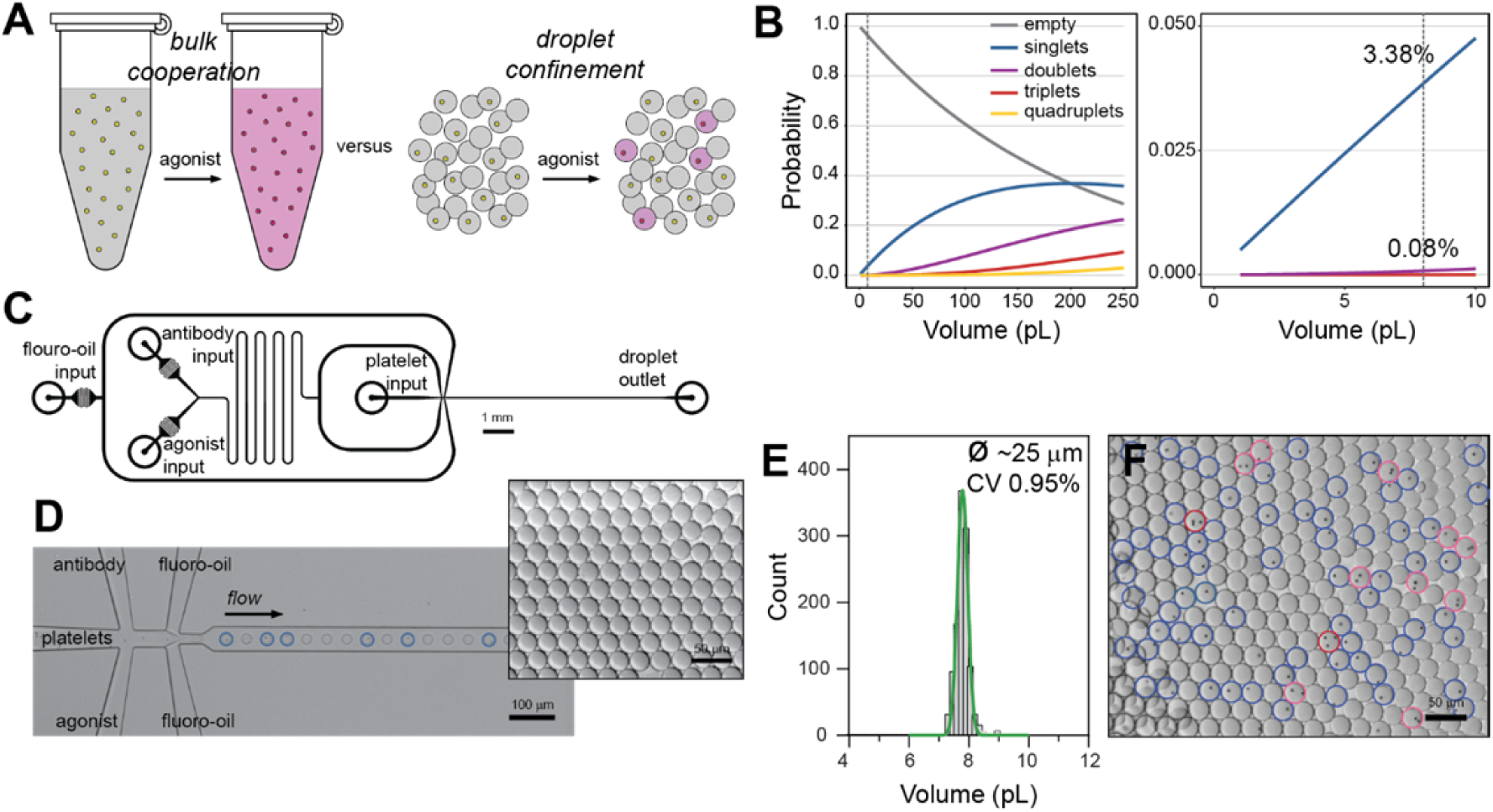
Concept and Methodology. Platelet populations cooperate during bulk perturbation experiments to produce an ensembled response whereas droplet compartmentalisation prohibits paracrine signalling to enable single platelet sensitivity measurements (A). The Poisson distribution is used to determine an optimal droplet volume for single platelet packaging with minimal multiple platelets (B). The microfluidic circuit for combining agonists and antibodies with platelets immediately before droplet generation is drawn to scale (3:1) (C). High throughput (10 kHz) droplet generation and single 2 μm particle packaging (blue circles) (D). Inset, droplet monodispersity is indicated by hexagonal packaging. Microfluidic conditions produce ∼8 pL (CV±0.95%) droplets (E). Poisson distribution impacting encapsulation illustrated using an excessive, 125 M/mL, 2 μm platelet-sized particle input concentration (F); single particle occupancy (blue; 23.9%), doublets (pink; 3.6%) and triplets (red; 0.7%).

To evaluate single platelet sensitivity differences a dose response experiment involving stimulating droplet-confined single platelets with convulxin (a GPVI receptor agonist) was undertaken and compared with the stimulation of platelet collectives. Using α_IIb_β_3_ activation (inside-out signalling) as the analytical end-point the platelet collectives produced a sigmoidal response curve emerging at 0.1 ng/mL and saturating at 1 ng/mL concentrations. The signal intensity distribution of the collective population indicates normally distributed functional variety. In comparison, a higher activation threshold is evident with singularly stimulated platelets, with activation emerging at 1 ng/mL and saturating at 10 ng/mL levels (Figure 2A). Extending the analysis to a different pathway, the P-selectin exposure end-point for alpha granule secretion, the same increased activation threshold for singularly stimulated platelets was observed (Figure 2B). Activation and aggregation density plots for platelets stimulated at 3 ng/mL are shown in Figure 2C and shows the hypersensitive behaviour of the collective response, the correlation between the two end-points and the bimodal distribution for singularly stimulated platelets undergoing population-level transition. Importantly, the hypersensitive sub-population was not observed by platelet collective dilution (up to a further 100-fold dilution), demonstrating the merit of the droplet microfluidics approach for single platelet analysis. To measure the significance in the response differences the relative risk was considered (Figure 2D,E). At low and high agonist concentrations the relative risk score is insignificant at 1.0, and at 3 ng/mL rises to 53 for α_IIb_β_3_ activation and 6 for P-selectin exposure end-points, highlighting the significantly (p<5×10^−5^) distinct hypersensitivity of collectively stimulated platelets.

**Figure 2.**
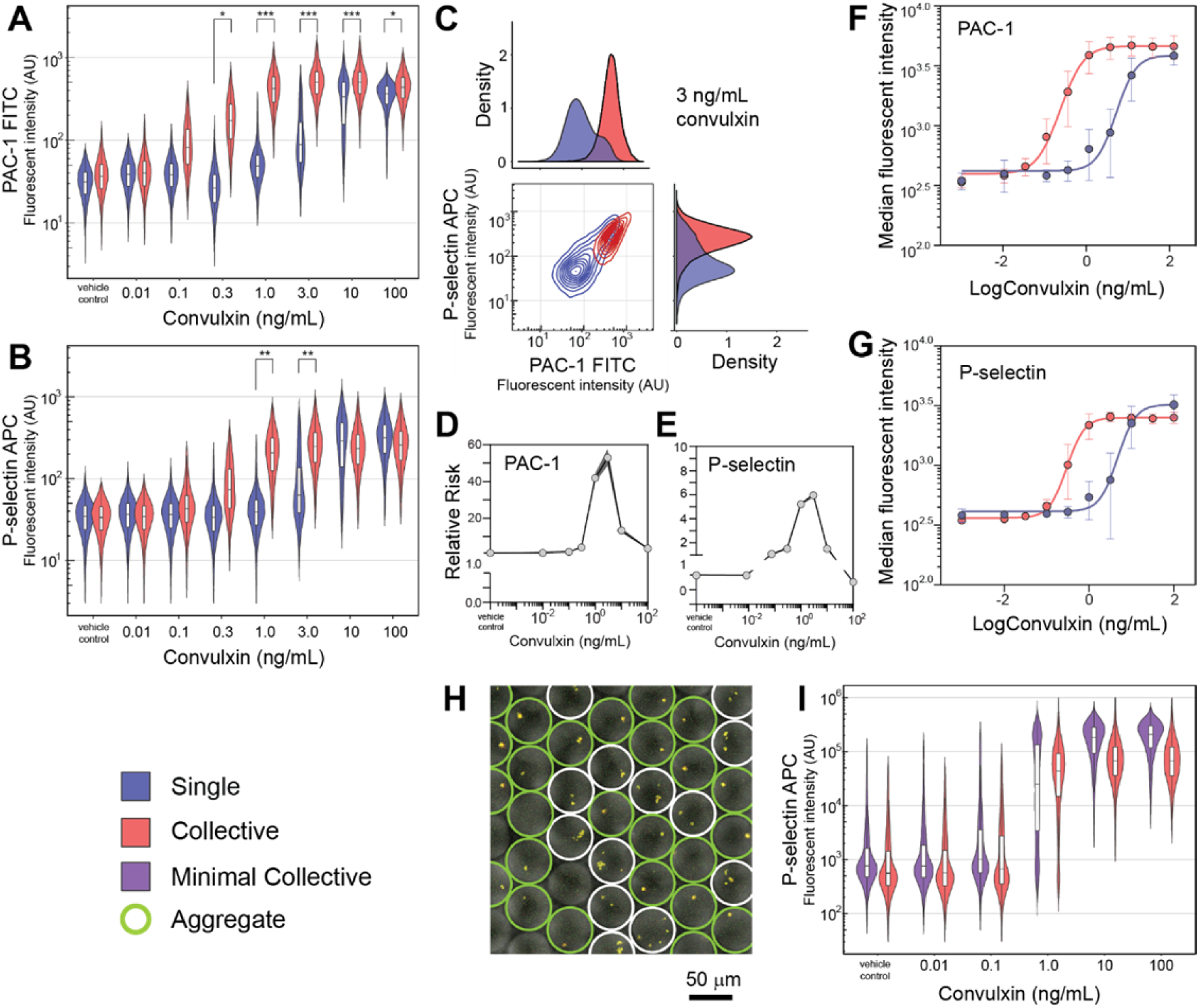
Broad-spectrum response to convulxin stimulation and hypersensitive collective behaviour. Violin plots comparing the activation of single platelets with platelet collectives using a convulxin dose response experiment, with PAC-1 binding to activated α_IIb_β_3_ (A) and P-selectin exposure (B) end-points. Cytometry plot and density plots of the emergence of hypersensitive single platelets at 3 ng/mL convulxin concentrations, while the collective population is fully activated (C). Relative risk analysis was used to determine the significance of the ∼20-fold differences between the single and collective platelet responses using PAC-1 (D) and P-selectin (E) end-points with confidence intervals determined by the Koopman asymptotic score. The *E*_*max*_ model was used to show a consistent difference between single and collective platelet behaviour across a diverse cohort (age; gender; smoking; BMI; exercise) of healthy donors using PAC-1 (F) and P-selectin (G) end-points. Droplet volume scaling to 65 pL produces minimal collectives (0–15 platelets with ∼5×10^8^ platelet/mL inputs) to allow aggregation responses to be investigated. Dual fluorescent imaging (P-selectin and CD63) with brightfield overlay of minimal platelet collectives stimulated with 1 ng/mL convulxin (H) and dose response violin plots of minimal platelet collectives compared with bulk platelet collective responses (I).

The sensitivity gains emerging from collective platelet behaviour were reproducible, with equivalent dose responses, both single and collective, obtained from the same donor 3 times over a 9 month period (**Supplementary Figure 2**). When the study was extended to a cohort of 8 healthy yet diverse donors (gender, age, BMI, smoking) the same pattern was observed, confirming the generality of the hypersensitive collective response, and allowing an efficacy model to be generated. For both α_IIb_β_3_ activation and P-selectin end-points, the collective convulxin response had an EC50 value of 0.4 ng/mL, whereas the single platelet EC50 was 7.5 ng/mL (Figure 2F,G). The 19-fold median sensitivity gains demonstrates the importance of hypersensitivity platelets and their cooperative influence.

To confirm that the molecular α_IIb_β_3_ activation and P-selectin end-points represent functional behaviour the dose response study was extended to larger droplets (65 pL; ø50 μm) packaging 0–15 platelets. At low concentrations (0.01 ng/mL) platelets are observed as discrete entities, whereas with moderate concentrations (1 ng/mL) single platelet aggregates are observed in droplets containing hypersensitive platelets, and not in droplets without hypersensitive platelets. At maximal concentrations (100 ng/mL) all droplets contain platelet aggregates (Figure 2(H)). Plotting the cytometry data shows a closer similarity with the collective platelet response (Figure 2(I)). However, a distinct bimodal distribution still results using 1.0 ng/mL convulxin. Elevated P-selectin signals relative to bulk conditions are also observed at 10 and 100 ng/mL convulxin. This is indicative of autocrine signalling resulting from the accumulation of degranulation products within the droplets. This experiment demonstrates the functional consequence of broad-spectrum sensitivity with cooperation and also that minimalistic platelet cooperation models can be used to understand transition states and the linkage between probabilistic molecular events and collective functional outcomes.

Collective sensitivity gains are attributed to the existence of low abundance hypersensitive platelets which, upon activation, degranulate to activate platelets in the vicinity that were insensitive to the initial stimulus. These modes of paracrine signalling produce a spatiotemporal corralling effect that drives platelet cooperation to deliver the collective response. Nevertheless, sufficient numbers of activated platelets are required to polarise the entire platelet population into an activated response (*e.g.* Figure 1A; collectives with 0.1 ng/mL convulxin). Our experiment involved platelets diluted to approximately 1/100^th^ of *in vivo* concentrations, suggesting digital activation may well occur under physiological conditions with insufficient volume to disperse paracrine signals. Platelet cooperation is mediated through the secretion of alpha granules as evidenced by P-selectin exposure, but also ADP and serotonin secretion from dense granules as evidenced by CD63 presentation (**Supplementary Figure 3**). The dense granule secretion pathway has a higher activation threshold than the alpha granule secretion pathway, consistent with the behaviour of these weaker agonists which augment the activation of other pathways for specialized platelet activation^51-53^. Again, autocrine signalling wherein stimulatory molecules accumulate in the droplets results in enhanced activation (α_IIb_β_3_ activation). This is evident with activation transition at 3 ng/mL convulxin stimulation, producing a clear bimodal distribution. In comparison, platelet collectives undergoing activation transition (0.3 ng/mL) have a lower α_IIb_β_3_ activation signal maxima than the droplet-confined hypersensitive single platelet sub-population (**Supplementary Figure 4**).

The study was extended to other agonists; the peptide TRAP-14 functional motif was used in place of thrombin to activate the PAR-1 receptor and as before α_IIb_β_3_ activation and P-selectin aggregation end-points were measured. The median activation threshold was again increased for single platelets stimulated in droplets, indicating that coordination by rare hypersensitive platelets reduces the activation threshold for platelet collectives. The emergence of a bimodal population distribution with singularly stimulated platelets was also observed for both end-points at 12.5 and 25 μM TRAP-14 concentrations (Figure 3A and 3B). The sigmoidal dose response again signifies continuous sensitivity variation. A small yet sensitive (∼4-fold) sub-population of single platelets stimulated with 12.5 μM TRAP-14 are evident. Experiments with the weak agonist ADP, stimulating the P2Y_1_ and P2Y_12_ receptors showed minor sensitivity gains, with a small but hypersensitive single platelet sub-population identified from droplets at low 0.1 μM ADP concentrations (**Supplementary Figure 5**).

**Figure 3.**
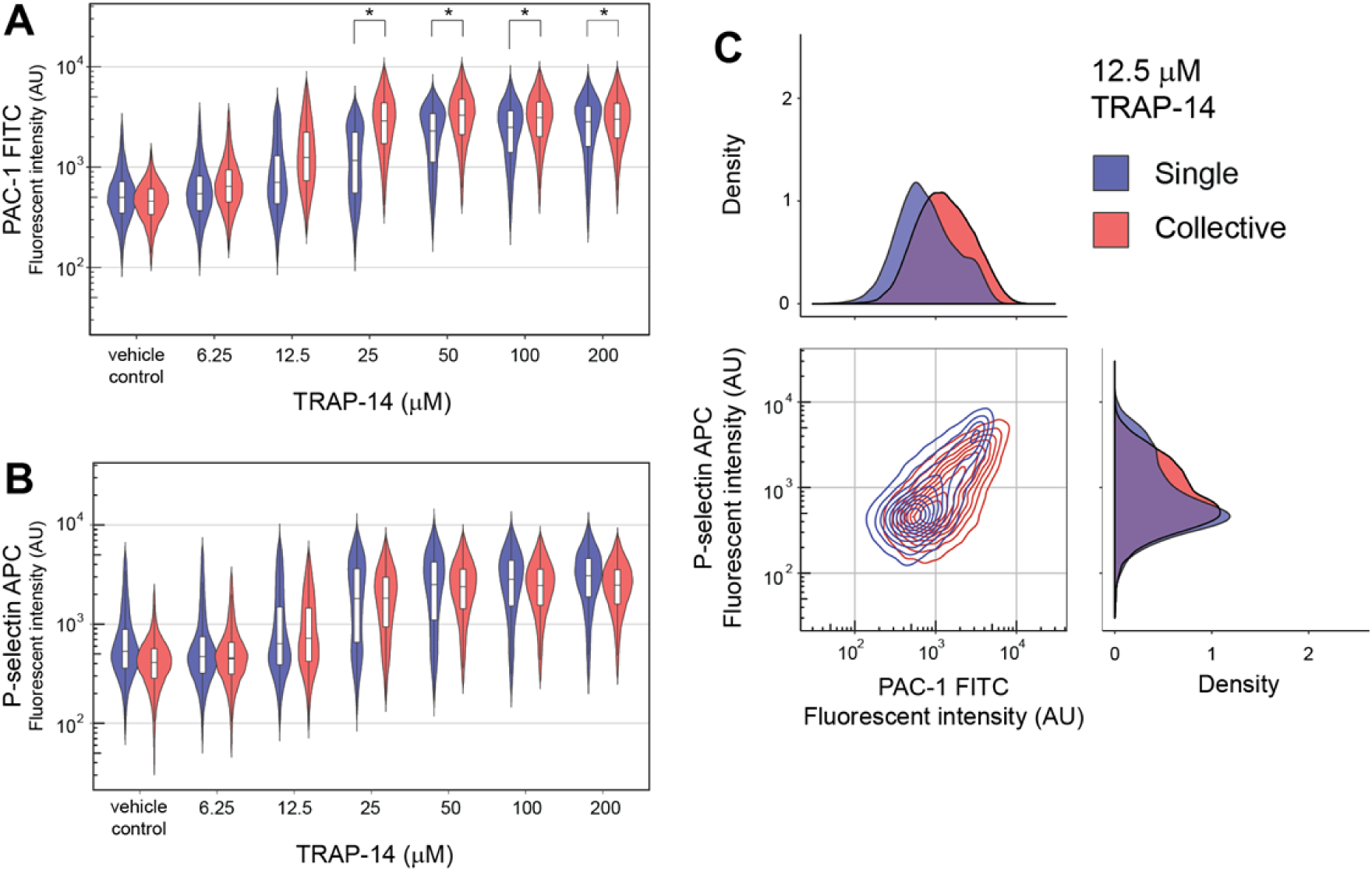
Variable TRAP-14 response and increased collective sensitivity. Violin plots comparing the activation of single platelets with platelet collectives using a TRAP-14 dose response experiment, with PAC-1 binding to activated α_IIb_β_3_ (A) and P-selectin (B) end-points. Cytometry and density plots showing increased collective sensitivity and the emergence of the more sensitive single platelet sub-population with a 12.5 μM TRAP-14 stimulation (C).

The collective sensitivity gains are agonist dependent, scaling with convulxin>TRAP-14>ADP and correlating with the potency of the agonist^19^. This scaling correlates with the steps in the clotting cascade, from platelet recruitment by collagen (≈convulxin) exposure, then thrombin produced by the coagulation cascade and lastly ADP secretions from the dense bodies of activated platelets in the vicinity. This demonstrates that droplet confinement does not downregulate platelet activation and implies collective sensitivity gains are most advantageous for triggering thrombus formation upon stimulation with collagen. Overall, collective sensitivity gains represents a strategy for robust, consensus-level, homeostasis emerging from paracrine cooperativity. This also conveniently exploits inherent platelet variability, thereby bypassing the need for functionally uniform platelets. Whether this diversity model involving community cross-talk for the transition from a dispersed state to localised recruitment and responsiveness can be generalised to other scenarios such as immune infiltration remains to be seen.

Functional variety is a common feature of cellular systems enabling powerful system responsiveness and control. This research shows that broad and continuous sensitivity distributions of single platelets interfaced via paracrine signalling produces robust collective sensitivity gains. During dual stimulation with collagen and thrombin, suspension phase platelets are known to polarise into two distinct populations; pro-coagulant and pro-aggregatory phenotypes^25^. The intrinsic or extrinsic nature of this heterogeneity is a matter of debate^8^. Again using droplet confinement we sought to resolve this debate and also to question the role of collective hypersensitivity in the emergence of the heterotypic response.

A dual stimulation dose response experiment was undertaken, with the responses of single and collective platelet populations compared using violin plots (**Supplementary Figure 6**). With platelet collectives pro-coagulant (annexin V high; α_IIb_β_3_ low) and pro-aggregatory (annexin V low; α_IIb_β_3_ high) heterotypic states emerged with a 100 ng/mL convulxin and 1.0 U/mL thrombin stimulation and is consistent with the literature.^16,17,23,24^ At the same concentrations droplet-confined, single platelet populations do not fully polarise, with an unresponsive third population (Figure 4A). Again this demonstrates the need for cooperation to enhance system sensitivity to activate all platelets. At higher dual stimulation doses (300 ng/mL convulxin and 3.0 U/mL thrombin) single platelet populations fully polarise into pro-coagulant and pro-aggregatory states. By excluding paracrine cross-talk, this confirms the intrinsic origins of heterogeneity. Indeed, removal of paracrine cooperative effects produces a fully digital pro-coagulant or pro-aggregatory response (Figure 4B). Importantly, these findings are made possible by single platelet confinement, advocating the use of droplet microfluidics to accurately delineate intrinsic single platelet phenotypes. In contrast to droplet-confined stimulation, the heterotypic distribution of platelet collectives involves some platelets with graded intermediate states. This implies the role of extrinsic effects for the generation of more subtle phenotypes likely required to enable more sophisticated functionality throughout the thrombus.

**Figure 4.**
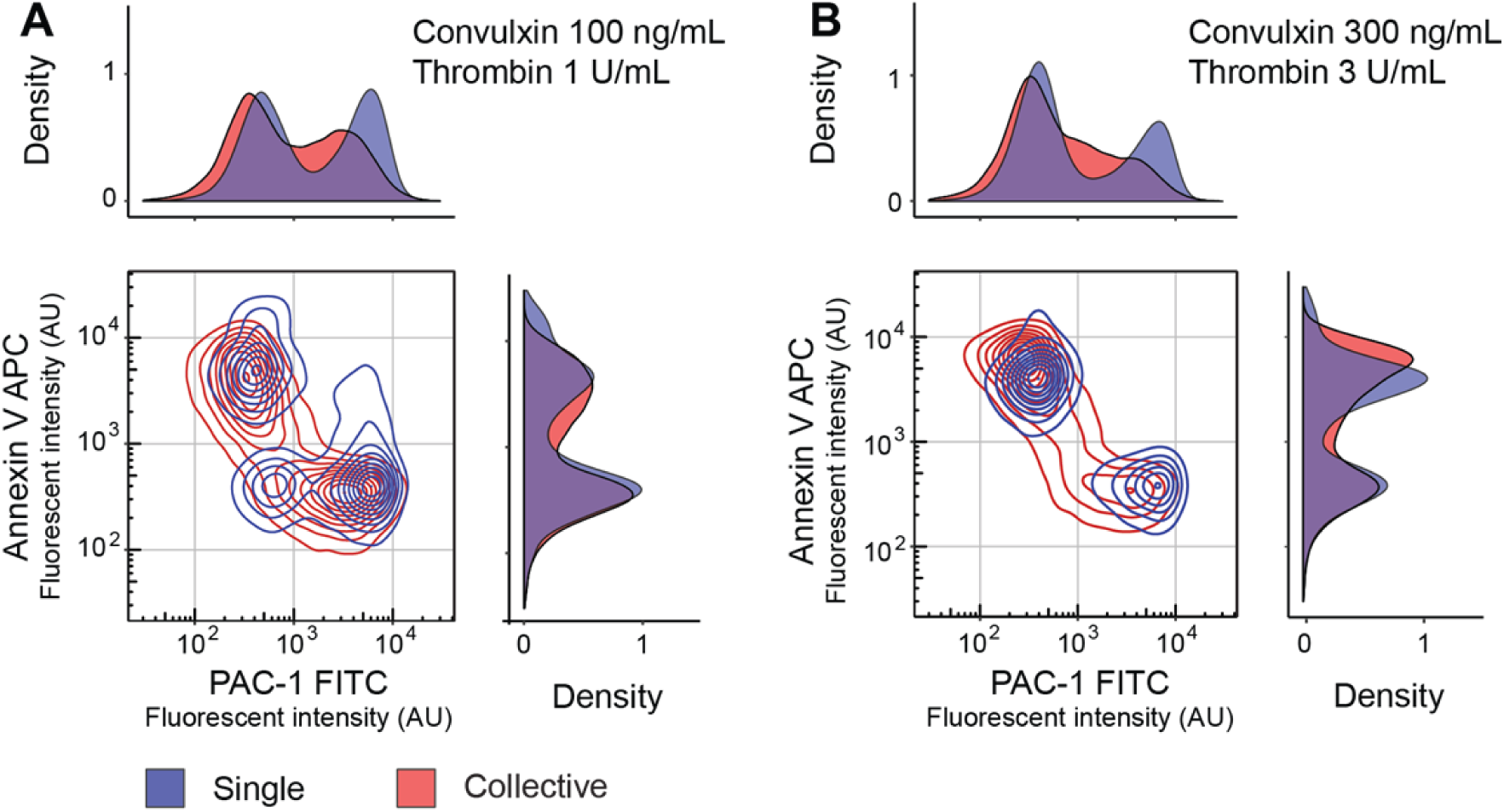
Intrinsic heterotypic states in response to dual stimulation. Stimulation of platelet collectives with 100 ng/mL convulxin and 1 U/mL thrombin produces pro-coagulant (annexin V high; α_IIb_β_3_ low) and pro-aggregatory (annexin V low; α_IIb_β_3_ high) states. With the same stimulation, single platelets produce a third unresponsive population (annexin V low; α_IIb_β_3_ low), indicating the requirement for paracrine cooperation to achieve complete population activation (A). Single platelets stimulated with higher 300 ng/mL convulxin and 3 U/mL thrombin concentrations drives platelets exclusively to functionally distinct pro-coagulant or pro-aggregatory states (B). Cooperation in platelet collectives at both dual stimulations concentrations directs some platelets into graded, intermediate activation states.

In this study, collective sensitivity gains are shown to be a general feature of human platelet biology. To gain further insights into this behaviour, these platelets were gated for characterisation (**Supplementary Figure 7**). Their forward and side scatter properties are indistinguishable from other, insensitive platelets. The CD42b signal (monomer component of the Von Willebrand factor receptor, GPIb-IX-V) for the hypersensitive platelets is similar albeit slightly reduced as a consequence of matrix metalloproteinase excision upon activation^54^. Further investigations involving large-scale antibody panels for highly multiplexed cytometry or more global proteomic^55^ and even transcriptomic screens^56,57^ following platelet sorting will be needed to determine the composition of the hypersensitive platelet sub-population.

Taken together our results show that collective platelet dynamics are dependent upon a hypersensitive subpopulation. In principle, such hypersensitivity could arise from natural variation within a functionally homogeneous platelet population or may be an important characteristic of a functionally distinct subpopulation. In the former case, all platelets are essentially equivalent in their functional responses, yet if platelet activation is an inherently stochastic process then the time-to-activation of each individual platelet will be described by a random variable. In this case, the observed variation in response may arise as a consequence of this underlying temporal stochasticity rather than being due to functional variability in the population per se. Similar stochastic mechanisms have been shown to be important in generating functional heterogeneity in other contexts^58^, for example within stem cell populations^59^. Alternatively, hypersensitive platelets may comprise a genuinely functionally distinct subpopulation. Similar issues have been seen in other biological contexts. For example, in the classic case of the emergence of bacterial resistance to virus infection, Luria and Delbrück applied a fractionation methodology with modelling to determine that stochastically acquired mutations, not a pre-existing subpopulation, produced resistance^60^. We anticipate that such a combination of experimental precision with single cell resolution and mathematical models^61^ will help resolve this issue.

## Conclusions

In this research we have developed a high throughput droplet microfluidics and flow cytometry methodology for measuring single platelet phenotypic and functional variability. The methodology was used to identify a broad-scale sensitivity continuum containing hypersensitive platelets which coordinate collective sensitivity gains by paracrine cooperativity to produce a robust system response. This feature can drive system polarisation into pro-aggregratory and pro-coagulant states during dual stimulation. Imbalance in the platelet population structure represents a potential route to pathology, either bleeding or arterial thrombosis leading to heart attacks and strokes. This discovery methodology can be used to characterise the nature of hypersensitive platelets and also has the potential to identify system or platelet population level prognostic biomarkers and potentially new therapeutic targets and intervention mechanisms.

## Materials and Methods

### Device Design and Fabrication

Microfluidic channels were 20 μm in height and with a width of 22 μm at the droplet generation junction for the reproducible generation of 25 μm (∼8 pL) droplets. The complete device design is available in the supplementary information (SI CAD), and involves separate agonist and antibody inlets that combine with the platelet inlet in advance of interfacing with the fluoro-oil phase at the droplet generation junction. All inlets, excepting the platelet inlet, included filter structures to remove any particulate and fibre contaminants during droplet formation. Microfluidic devices were prepared by standard SU-8 photolithography followed by poly(dimethylsiloxane) (PDMS, Sylgard 184) to polyurethane (Smooth-On 310) mould cloning for parallel replication by soft lithography in PDMS at 60°C for 2 hours. Inlet/outlet ports for plug and play interconnection were produced using a 1-mm-diameter Miltex biopsy punch (Williams Medical Supplies Ltd). Devices were bonded to PDMS-coated glass microscope slides using a 30 s oxygen plasma treatment (Femto, Deiner Electronic) followed by surface functionalisation using 1% (v/v) trichloro(1*H*,1*H*,2*H*,2*H*-perfluorooctyl)silane (Sigma Aldrich) in HFE-7500™ (3M™ Novec™). Minimal platelet collectives were encapsulated in 50-μm-diameter droplets, generated with a 50-μm high microfluidic device with a 50-μm-wide droplet generation junction that were fabricated as described above.

### Participants and Sampling

Blood was obtained from healthy volunteers after obtaining written consent (REC: 14/SC/0211). Participants were free from anti-platelet medication, such as aspirin for 2 weeks and >24 hours free from other non-steroidal anti-inflammatory medication. The cohort was diverse, with 5 male and 3 female volunteers, aged between 20 and 60 and with one smoker. Venepuncture with a 21G needle was used to collect blood in vacuum tubes containing 1:10 v/v 3.2% trisodium citrate (first 4 mL discarded). Platelet counts were determined using the method described by Masters and Harrison^62^, involving a CD61 antibody and an Accuri C6 instrument (BD Biosciences). These tubes were gently inverted, centrifuged at 240 g for 15 minutes without brake to prepare platelet rich plasma (PRP) that was rested for 30 minutes prior to experiments, and diluted to a concentration of 25×10^6^/mL in HEPES buffer (136 mM NaCl, 2.7 mM KCl, 10 mM HEPES, 2 mM MgCl_2_, 0.1% (w/v) glucose and 1% (w/v) BSA (pH 7.45)) for dose response experiments.

### Droplet Microfluidics

Medical grade, sterile polythene tubing (ID 0.38 mm; OD 1.09 mm) was used to directly interface syringes with 25G needles to the microfluidic ports. Syringe pumps (Fusion 200, Chemyx) were used to deliver reagents. The Poisson distribution effect was evaluated using NIST, monodisperse 2-μm-diameter polystyrene particles (4202A, ThermoScientific™). Platelet experiments involved the delivery of HFE-7500 fluoro-oil (3M™ Novec™) with 0.75% (v/v) 008-fluorosurfactant (Ran Biotechnologies) at 20 μL/min, antibody and agonist solutions at 2 μL/min and platelets at 1 μL/min to generate 25-μm-diameter droplets. High speed imaging (2,500 fps) using a Miro eX2 camera (Phantom) mounted on an open instrumentation microscope (dropletkitchen.github.io) was used to document droplet generation and an inverted fluorescent microscope (CKX41, Olympus) fitted with a QIClick camera (Teledyne, QImaging) was used to image droplet contents. Droplets were collected for 5 minutes, incubated while resting at room temperature in the dark for 10 minutes, then combined with CellFix fixative (BD Biosciences) and subsequently with 1*H*,1*H*,2*H*,2*H*-perfluoro-1-octanol (PFO, Sigma-Aldrich) to destabilise the droplet interface and break the emulsion. Control experiments involved the same agonist and antibody treatments of platelet collectives in microcentrifuge tubes. In the case of the 50 μm droplets, the reagent flow rates were: 80 μL/min for fluoro-oil, 4 μL/min undiluted platelet rich plasma (∼5×10^8^/mL), and 8 μL/min for convulxin and antibody inputs. Platelets were incubated for 60 minutes prior to emulsion breaking and fixation.

### Flow Cytometry

Platelets were stimulated with convulxin (Enzo Life Sciences), a snake venom which activates the GPVI receptor, TRAP-14 (Bachem AG) an agonist of the PAR-1 receptor or ADP (Sigma Aldrich) an agonist of the P2Y_12_ receptor. The dual agonist experiment involved stimulation with convulxin and thrombin (Sigma) in the presence of 2.5 mM CaCl_2_. Here, coagulation was prevented using 0.5 μM rivaroxaban (Advanced ChemBlocks Inc) and 100 mM H-Gly-Pro-Arg-Pro-OH (GPRP, Bachem) added to the HEPES platelet dilution buffer. Fluorescent antibodies and selective stains were used to detect biomarkers: Fluorescein isothiocyanate (FITC) conjugated PAC-1 (PAC-1 clone at 1.25 ng/μL), allophycocyanin (APC) conjugated CD62P (P-selectin, AK-4 clone at 0.63 ng/μL), FITC conjugated anti-CD63 (H5C6 clone at 2.0 ng/μL), R-phycoerythrin (PE) conjugated CD42b (HIP1 clone at 1.25 ng/μL) and Annexin V at 0.08 ng/μL were obtained from BD Biosciences. Following treatments, antibody incubation and fixation, samples were diluted in PBS and measured using an Accuri C6 flow cytometer (BD Biosciences). Platelets were identified using a gate on CD42b-PE intensity, with doublets and non-platelet-sized events removed by gating. In the case of the 50 μm droplets, CD42b-PE, CD63-FITC and CD62P-APC antibodies were used and incubated for 60 minutes prior to fixation and emulsion breaking.

### Statistics

Droplet images were analysed using ImageJ (NIH) and flow cytometry data using FlowJo. To compare single and collective platelet responses from a single donor, the relative risk statistic was used to quantify the association between stimulation and response (R, epitools). The overall cohort response difference between single and collective platelets was plotted using an efficacy maxima (*E*_*max*_) sigmoidal model generated in GraphPad Prism.

## Acknowledgements

The research was funded by the Marie Curie (333721, JW), the British Heart Foundation (FS/13/67/30473, MSAJ) and the Medical Research Council (MC_PC_15078, MSAJ). We thank Simon Lane for support with ImageJ analysis, Johan WM Heemskerk for critical manuscript feedback and Joanna D Stewart for proofing the manuscript.

## Author contributions

J.W. conceived the project, M.S.A.J. did the experiments and analysed the data, N.A.E., B.D.M. and J.W. supervised the research, J.W. wrote the manuscript and M.S.A.J., N.A.E. and B.D.M. reviewed the manuscript and approved the final version.

## Declarations

The authors declare no competing interests.

## Supporting Information

Droplet microfluidic design file (SI_CAD.dwg).

