## Supplementary Figures and Data for "Single platelet variability governs population sensitivity and initiates intrinsic heterotypic behaviours"

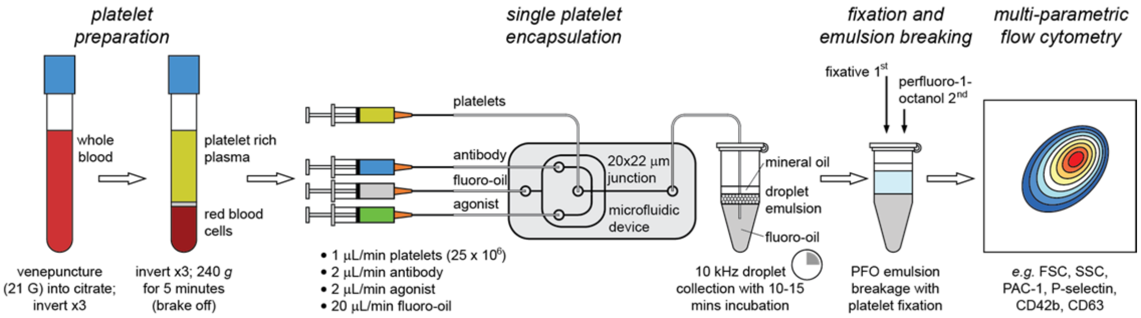


Supplementary Figure 1. *Illustrated protocol.* Venous blood draw through to platelet isolation, droplet encapsulation, incubation, platelet release by breaking the emulsion with fixation in readiness for flow cytometry. The analytical pipeline involves kHz droplet generation and kHz flow cytometry to deliver the throughput necessary to comprehensively interpret platelet population functional distributions.

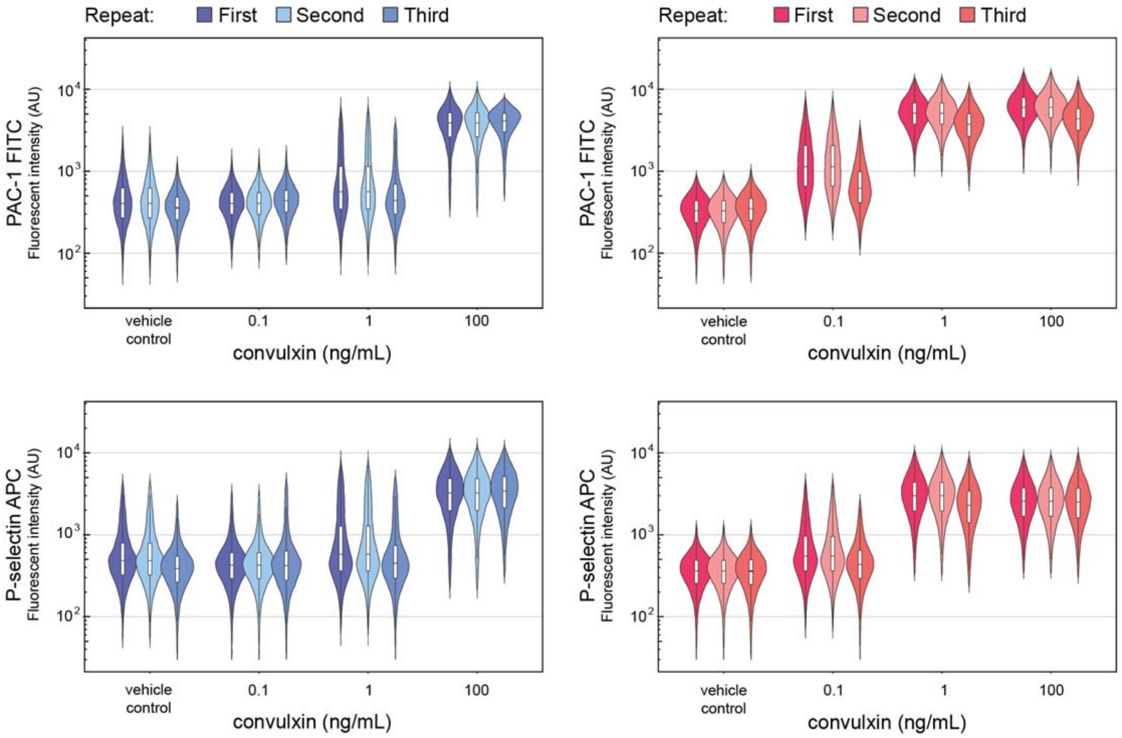


Supplementary Figure 2. *Donor reproducibility.* PAC-1 (top) and P-selectin (bottom) single (blues) and collective (reds) platelet responses to convulxin stimulation for the same donor measured three times over a 9 month period.

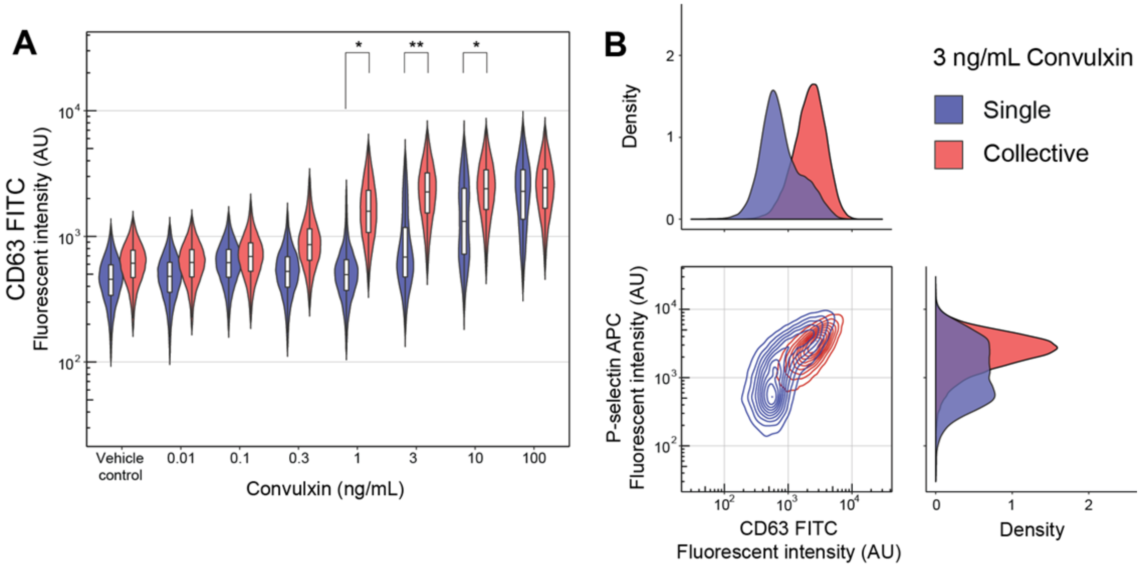


Supplementary Figure 3. *Alpha and dense granule secretions correlate.* Dense granule (CD63) secretion is consistent with the alpha granule response (P-selectin) for single platelets stimulated in droplets with 3 ng/mL convulxin. However, the dense granule secretion pathway has a higher threshold for complete activation.

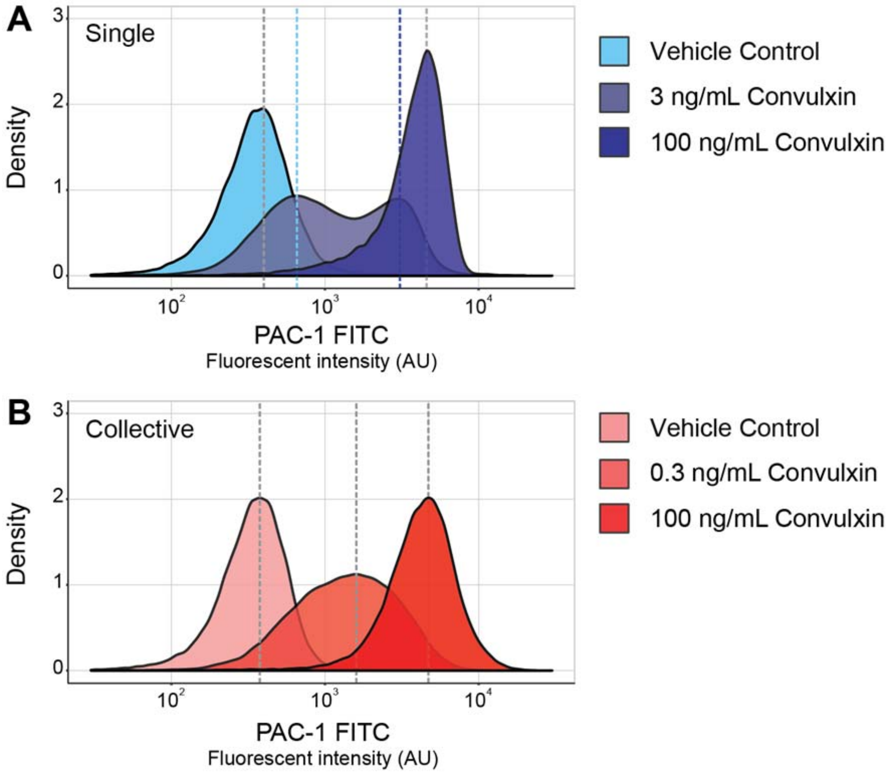


Supplementary Figure 4. *Transition states differ between single and collective platelet populations.* The peak maxima for inactive, transition and active states are identified with dashed lines for singular (A) and collective (B) platelet populations. The bimodal single platelet transition involves distinct inactive (light blue) and active (dark blue) signal maximas, the latter higher than the maxima for collective platelets undergoing transition. This is attributed to autocrine feedback during droplet confinement.

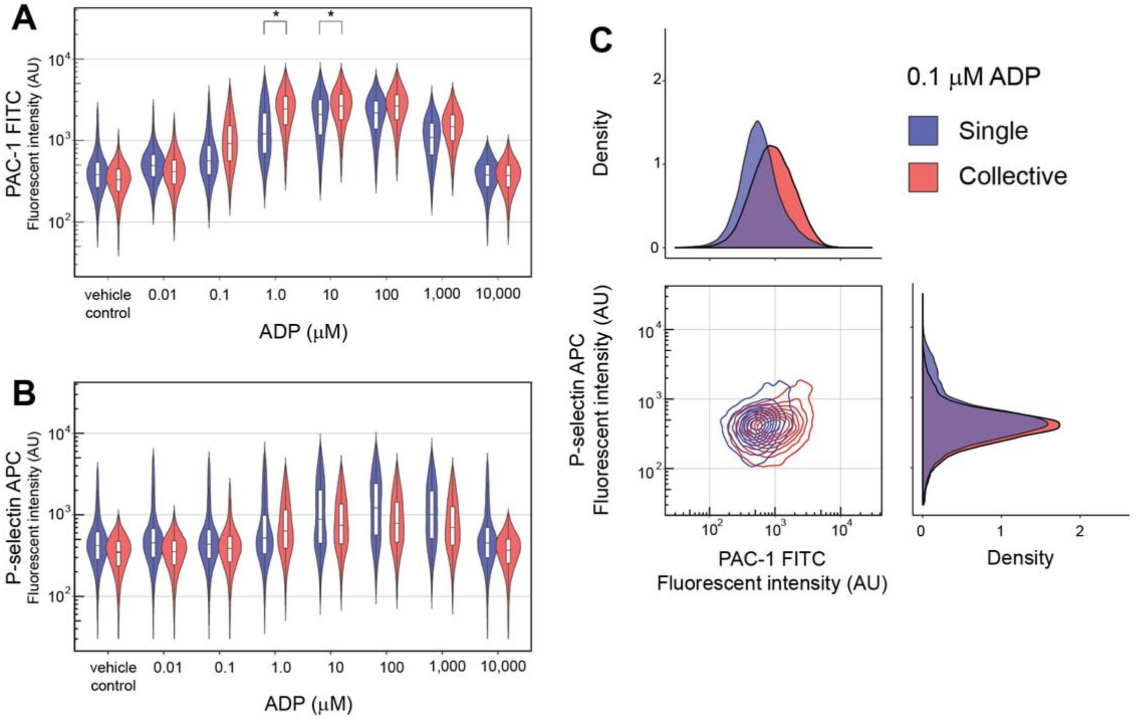


Supplementary Figure 5. *Minor variability in the response to ADP and minor collective sensitivity gains.* Violin plots comparing the activation of single platelets with platelet collectives using an ADP dose response experiment, with PAC-1 binding to activated $\alpha_{IIb}\beta_3$ (A) and P-selectin exposure (B) end-points. Cytometry and density plots showing a small yet hypersensitive subpopulation for single platelets in droplets at 0.1 μM ADP (C).

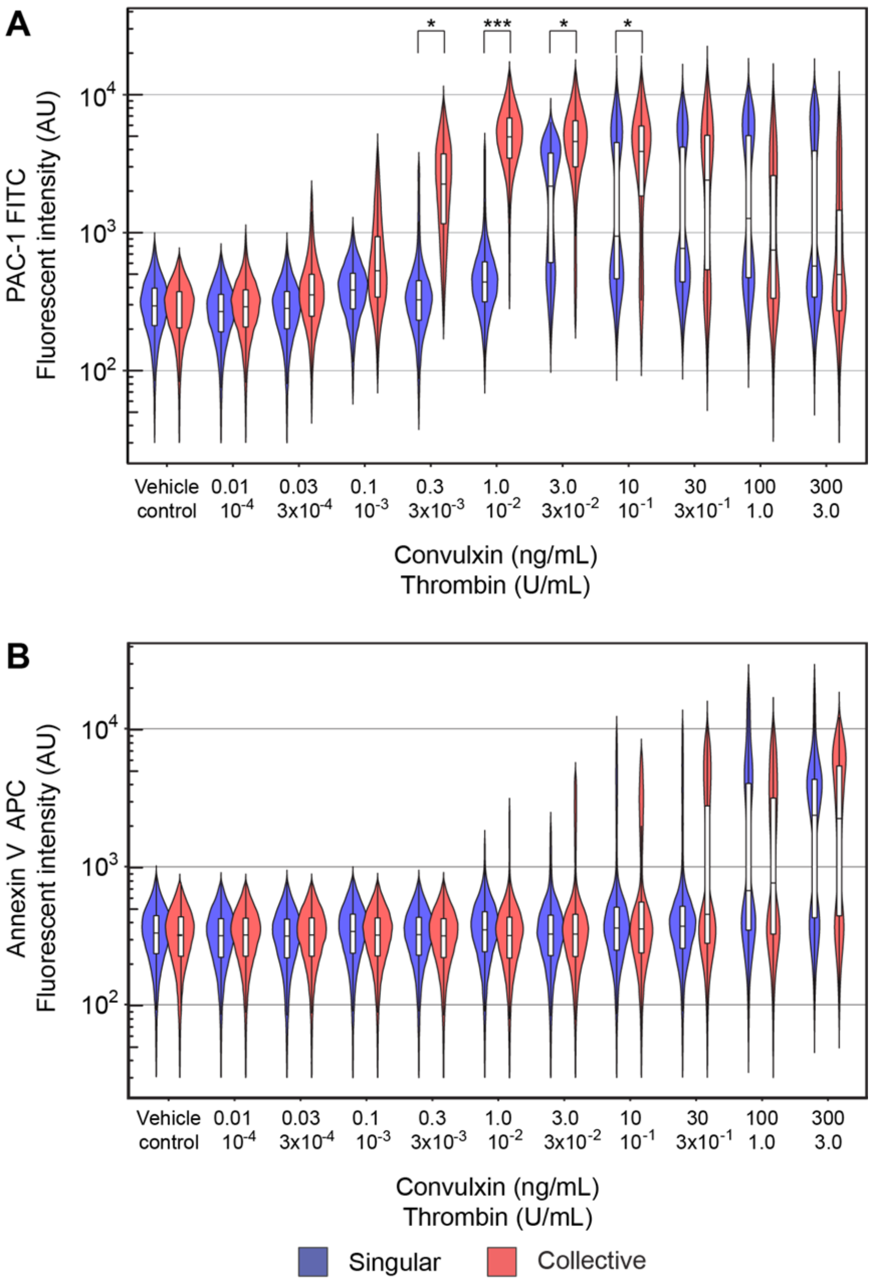


Supplementary Figure 6. *Dose responses for dual stimulated single and collective platelets.* PAC-1 binding (A) and P-selectin exposure (B) were used as end-points.

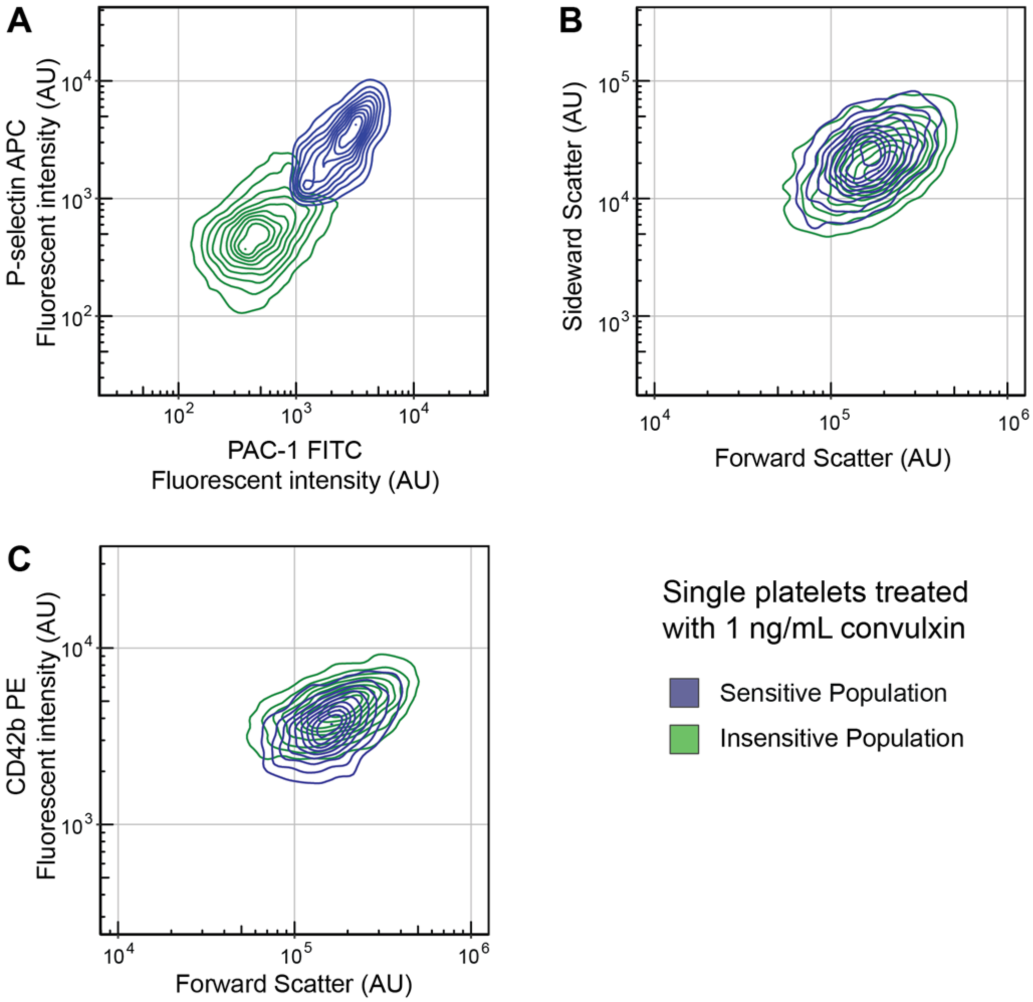


Supplementary Figure 7. *The nature of hypersensitivity requires further investigation.* Hypersensitive platelets ('sensitive population') identified by PAC-1 and P-selectin staining (A) cannot be distinguished by forward and side scatter properties from other ('insensitive') platelets (B). Platelet activation causes a minor CD42b signal reduction (C).
